# HDAC11 Deficiency Prevents High-Fat Diet-Induced Obesity and Metabolic Syndrome

**DOI:** 10.1101/310383

**Authors:** Lei Sun, Caralina Marin De Evsikova, Ka Bian, Alexandra Achille, Elphine Telles, Edward Seto

**Affiliations:** George Washington University Cancer Center, George Washington University School of Medicine & Health Sciences, Washington, DC 20037, USA; Department of Biochemistry & Molecular Medicine, George Washington University School of Medicine & Health Sciences, Washington, DC 20037, USA; Department of Molecular Medicine, Morsani College of Medicine, University of South Florida, Tampa, FL 33612, USA; Moffitt Cancer Center, University of South Florida, Tampa, FL 33612, USA

**Keywords:** HDAC11, obesity, metabolic syndrome, hepatic steatosis, UCP1, CPT1, adiponectin

## Abstract

Obesity is a serious and widespread health problem which has become a growing concern in many societies. Most currently available weight-loss medications do not work for everyone, and the effects decline over time. Thus, there is an urgent need to identify new molecular targets to improve drug development for the treatment of obesity and obesity-related diseases. In this study, we discovered that the histone deacetylase 11 (HDAC11) enzyme is a key regulator of metabolism and obesity, and the absence of HDAC11 prevents obesity in mice. Our findings will facilitate the development of novel therapeutics to treat obesity by targeting HDAC11.

**Abstract:** Obesity and its associated metabolic syndromes are the consequence of susceptible genes and obesogenic environments. We report here that histone deacetylase 11 (HDAC11) plays a critical role in the development of obesity and in metabolic homeostasis. HDAC11 knockout mice display resistance to high-fat diet-induced obesity and associated syndromes by enhancing glucose tolerance and insulin sensitivity, attenuating hypercholesterolemia and hyperinsulinemia, and blocking hepatosteatosis and liver damage. Mechanistically, HDAC11 deficiency boosts energy expenditure through promoting thermogenic capacity, which attributes to the elevation of uncoupling protein 1 (UCP1) expression and activity in brown adipose tissue. Moreover, loss of HDAC11 stimulates mitochondrial oxidation, elevates plasma adiponectin, and activates the adiponectin-AdipoR-AMPK pathway in the liver, which may contribute to a reversal in hepatosteatosis. These findings establish HDAC11 as a key regulator of metabolism and indicate that HDAC11 inhibitors may hold promise for treating overweight and obesity-related diseases.

## Introduction

Histone deacetylases (HDACs) regulate a wide range of biological functions by removing acetyl and acyl groups from lysine residues on histones and non-histone proteins. There are two categories of HDACs, the Zn^2+^-dependent deacetylases (HDAC1-11) and the NAD+-dependent deacetylases (SIRT1-7). Based on sequence similarities, eighteen HDACs have been grouped into four classes (1). Class I HDACs (HDAC1, 2, 3 and 8) are similar to *Saccharomyces cerevisiae RPD3*. Class II (HDAC4, 5, 6, 7, 9, and 10) share a similar core sequence with yeast *HDA1*. SIRT1-7 belong to Class III, which are orthologs of yeast *SIR2* (silent information regulator 2). HDAC11 is the sole Class IV HDAC, which shares its highly conserved critical residues in the catalytic core regions with both class I and II HDACs.

Many studies have established that abnormal HDACs play a key role in human diseases including cancer, neurological diseases, metabolic/endocrine disorders, inflammatory diseases, immunological disorders, cardiovascular diseases, and pulmonary diseases. As a result, there is significant interest in targeting HDACs for therapeutic purposes. In the area of metabolic/endocrine disorders, there is increasing evidence that Class I, II and III HDACs are intimately involved in the regulation of metabolism, and dysregulation of these HDACs may lead to maladies such as obesity and type 2 diabetes.

For Class I HDACs, overexpression of HDAC1 blocks, whereas deleting HDAC1 significantly enhances, β-adrenergic activation-induced BAT-specific gene expression in brown adipocytes, suggesting that targeting HDAC1 may be beneficial in the prevention and treatment of obesity by enhancing BAT thermogenesis. (2). It has been reported that hepatic deletion of HDAC3 leads to lipid accumulation in the liver (3) and disrupts normal metabolic homeostasis (4). HDAC3 inhibition, however, improves glycemia and insulin secretion in obese diabetic rats (5). Treatment of mice with MS-275, an HDAC1/HDAC3 inhibitor, stimulates the functionality and oxidative potential of adipose tissue, improves glucose tolerance and ameliorates the metabolic profile in diet-induced obese mice (6). In a contrasting study, inhibition of HDAC8 causes insulin resistance (7).

For Class II HDACs, high HDAC5 and HDAC6 expression levels are required for adequate adipocyte function (8). HDAC9 gene deletion prevents the detrimental effects of chronic high-fat feeding on adipogenic differentiation, increases adiponectin expression, and enhances energy expenditure by promoting beige adipogenesis, thus leading to reduced body mass and improved metabolic homeostasis (9).

Sirtuins, the Class III histone deacetylase family, are regulators of metabolism, and like Class I and II HDACs, may also be therapeutic intervention targets. Hepatocyte-specific deletion of SIRT1 alters fatty acid metabolism and results in hepatic steatosis and inflammation (10). Ablation of SIRT6 in mice results in severe hypoglycemia, as well as liver steatosis (11). Additionally, while neural-specific deletion of SIRT6 in mice promoted diet-induced obesity and insulin resistance (12), SIRT6 overexpression protected against these effects (13).

Unlike Class I, II, and III HDACs, there is limited information on the role of HDAC11, the sole Class IV HDAC, in the regulation of metabolic pathways and almost nothing is known about its potential role in obesity. In fact, most studies on HDAC11 have focused on its function in immune and neuro systems, as well as in cancer development. In addition to its potential modest deacetylase activity, recent studies suggest that HDAC11 is a long chain fatty-acid deacylase (14). Similar to posttranslational lysine acetylation/deacetylation, protein acylation/deacylation is a mechanism of biological signaling and many fatty acylated proteins play key roles in regulating cellular structure and function. Therefore, we sought to determine if HDAC11 may also impact metabolic phenotypes and core metabolic pathways.

Here we report a novel key regulatory function of HDAC11 in metabolic homeostasis. HDAC11 deletion protected mice from high-fat diet (HFD)-induced obesity, insulin resistance and hepatic steatosis. These results uncovered the potential prospect of HDAC11 inhibitors in therapy of obesity and obesity-related diseases.

## Results

### HDAC11 knockout mice display resistance to HFD-induced obesity

To study the function of HDAC11 in metabolic homeostasis, we first assessed if the loss of HDAC11 had any effect on animal body weight maintained under HFD. HDAC11 knockout (KO) mice were generated with a targeted deletion of floxed exon 3 of *Hdac11*, and the KO was confirmed by genotyping mouse-tail DNA derived from wild-type (WT) and HDAC11 KO mice (Figs. S1A, B). Loss of HDAC11 protein expression was also verified by Western blot analysis of protein extracts from WT and HDAC11 KO mouse brains (Fig. S1C).

When male WT and HDAC11 KO mice were fed HFD (42% kcal from fat; ad libitum from 3-week-old), a significant difference in body weight between the two groups was observed beginning at about 10 weeks (Fig. 1A). Maintaining the animals on HFD for up to 30 weeks resulted in approximately 25% more body weight gain in WT mice compared to HDAC11 KO mice (Fig. 1B). Furthermore, the epididymal white adipose tissue (eWAT) weight was significantly reduced in HDAC11 KO mice compared to WT after 20 weeks feeding on HFD (Fig. 1C). Hematoxylin and eosin (H & E) stain showed that the average size of adipocytes was smaller in the eWAT of HDAC11 KO mice compared to the WT mice (Figs. 1D, E), reflecting a reduction of lipid accumulation in HDAC11 KO mice.

**Figure 1.**
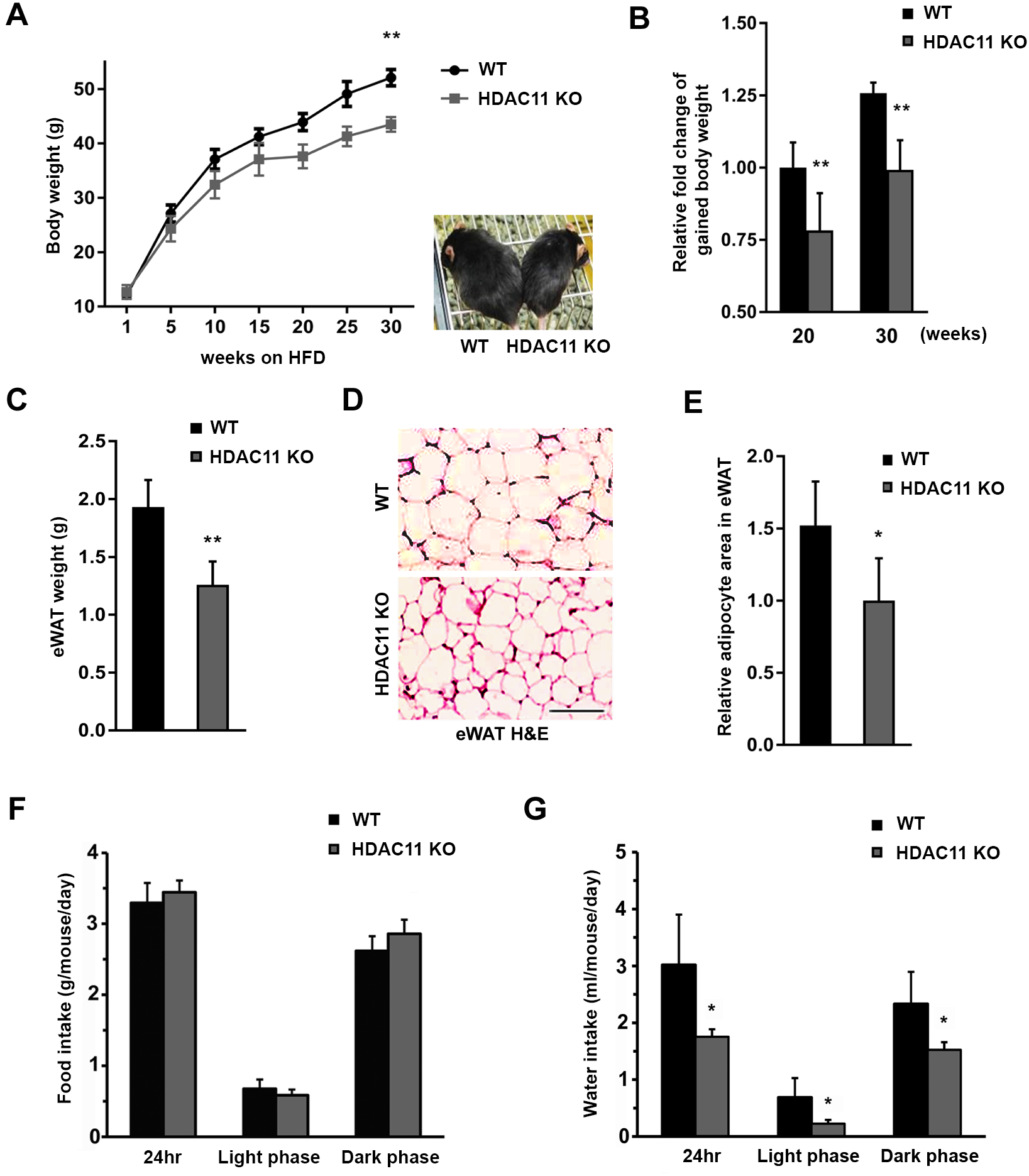
HDAC11 KO mice are resistant to HFD-induced obesity. (A) Body weights of WT (n=9) and HDAC11 KO (n=8) mice fed with HFD. A representative picture of a WT and a HDAC11 KO mouse after 20-weeks on HFD (right panel). (B) Comparison of body weight gain between WT and HDAC11 KO mice on 20-week and 30-week HFD. (C) Weight of epididymis white adipose tissue (eWAT) between WT (n=5) and HDAC11 KO (n=5) mice on 20-week HFD. (D) Representative H & E stain of eWAT sections, scale bar: 200 μm. (E) Relative adipocyte area quantified in the eWAT between WT and HDAC11 KO mice on 20-week HFD. (F) Food and (G) water intake averaged every day in WT (n=4) and HDAC11 KO (n=4) mice in 24 hours, light and dark phase. Data shown are mean ±SD.*p<0.05 and **p<0.01 compared to control group.

To investigate whether energy intake plays a role in HDAC11 KO-driven obesity resistance, we compared the amount of food consumption between the two groups using a comprehensive cage monitoring system for 96 hours. We found no statistical differences in food consumption between WT and HDAC11 KO mice (Fig. 1F). Further comparison of food intake for 17 consecutive weeks, showed that no marked difference was observed between WT and HDAC11 KO mice (Fig. S2). In contrast to food intake, water consumption was significantly increased in WT animals, and the higher water intake by WT mice occurred across both light phase and dark phase, suggesting a status of polydipsia (Fig. 1G).

### HDAC11 knockout mice are protected from HFD-associated hypercholesterolemia and hyperinsulinemia

To determine if HDAC11 affects hyperlipidemia under HFD, we tested cholesterol and triglyceride (TG) levels in plasma of WT and HDAC11 KO mice. HFD feeding for 20 weeks increased cholesterol level in WT mice, but not in HDAC11 KO mice (Fig. 2A). For plasma TG, there was a slight, but not significant, difference between HDAC11 KO and WT groups after 20-week of HFD (Fig. 2B), which is consistent with a previous report of decreased circulating TG levels in HFD fed C57BL/6 mice (15). We further measured liver TG level, which was markedly reduced in HFD-fed HDAC11 KO mice compared to WT mice (Fig. 2C).

**Figure 2.**
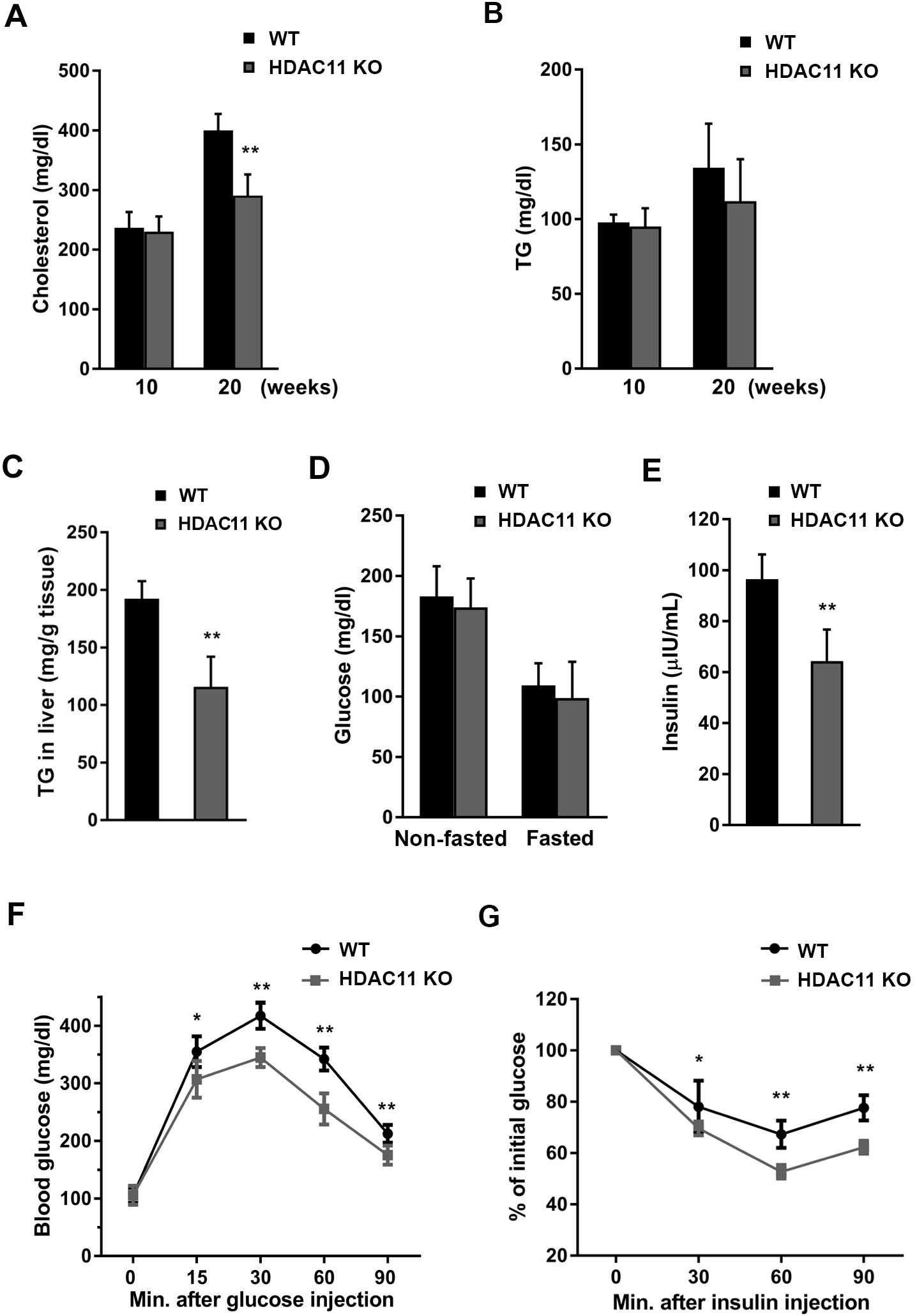
HDAC11 KO mice are protected from HFD-induced hypercholesterolemia and hyperinsulinemia. (A) Plasma cholesterol and (B) plasma triglyceride (TG) levels in WT (n=8) and HDAC11 KO (n=10) mice on 10-week and 20-week HFD. (C) Hepatic triglyceride (TG) content in WT (n=7) and HDAC11 KO (n=6) mice on 26-week HFD. (D) Plasma glucose level on 20-week HFD nonfasting and fasting for 12 hours in WT (n=8) and HDAC11 KO (n=10) mice. (E) Plasma insulin level on 20-week HFD in WT (n=8) and HDAC11 KO (n=10) mice. (F) Glucose tolerance test (GTT) and (G) insulin tolerance test (ITT) in WT (n=12) and HDAC11 KO (n=11) on 13-week HFD. Data shown are mean ±SD. *p<0.05, **p<0.01.

Because HDAC11 deficient mice have less excessive drinking behavior compared to WT mice, we measured both fasting and non-fasting blood glucose levels to assess the diabetes status in these animals. The results show that there is no marked difference between HDAC11 KO and WT mice (Fig. 2D). To further explore the role of HDAC11 in glucose metabolism, blood insulin levels were measured, which showed a significant decrease in HDAC11 KO mice (Fig. 2E). Also, compared to WT animals, HDAC11 KO mice show improved glucose tolerance as determined by the glucose tolerance test (GTT), which measures the clearance by the body of an injected glucose load (Fig. 2F). Similarly, using the insulin tolerance test (ITT), which monitors endogenous glucose disappearance over time in response to an insulin injection, HDAC11 KO mice have a significant greater reduction in blood glucose levels after insulin challenge compared to WT mice (Fig. 2G). Together these findings suggest that HDAC11 plays an important role in maintaining glucose homeostasis and insulin sensitivity. The effect of HDAC11 in HFD-associated hyperinsulinemia are not due to changes in the mRNA expression levels of insulin or glucagon receptors (Fig. S3). There is also no difference in the animals’ hematology examination results (Table S1).

### HDAC11 knockout mice are resistant to HFD-induced hepatic steatosis

Next, we examined obesity-associated hepatosteatosis and liver function. The livers of WT mice were paler and enlarged compared to HDAC11 KO mice after 26 weeks of HFD (Figs. 3A-C). Histological examination revealed severe steatosis and lipid accumulation in WT, but not in HDAC11 KO, mouse livers (Fig. 3C). Oil red O stain also showed extensive accumulation of large lipid droplets in the liver of WT, but not HDAC11 KO mice (Fig. 3D). These results are consistent with our observation that the liver TG level was significantly reduced in HFD-fed HDAC11 KO mice (Fig. 2C), and suggests a beneficial protective role of HDAC11 deletion in the HFD-associated hepatic steatosis. To further verify the role of HDAC11 on lipid accumulation in hepatocytes, we knocked-down HDAC11 expression (HDAC11 KD) in the mouse hepatic cell line AML12. In HDAC11 KD cells, lipid droplet accumulation significantly reduced after five days of treatment with free fatty acids, compared to control AML12 cells (Figs. 3E, F). Our results, therefore, strongly support a regulatory effect of HDAC11 on lipid accumulation and HFD-associated hepatic steatosis.

**Figure 3.**
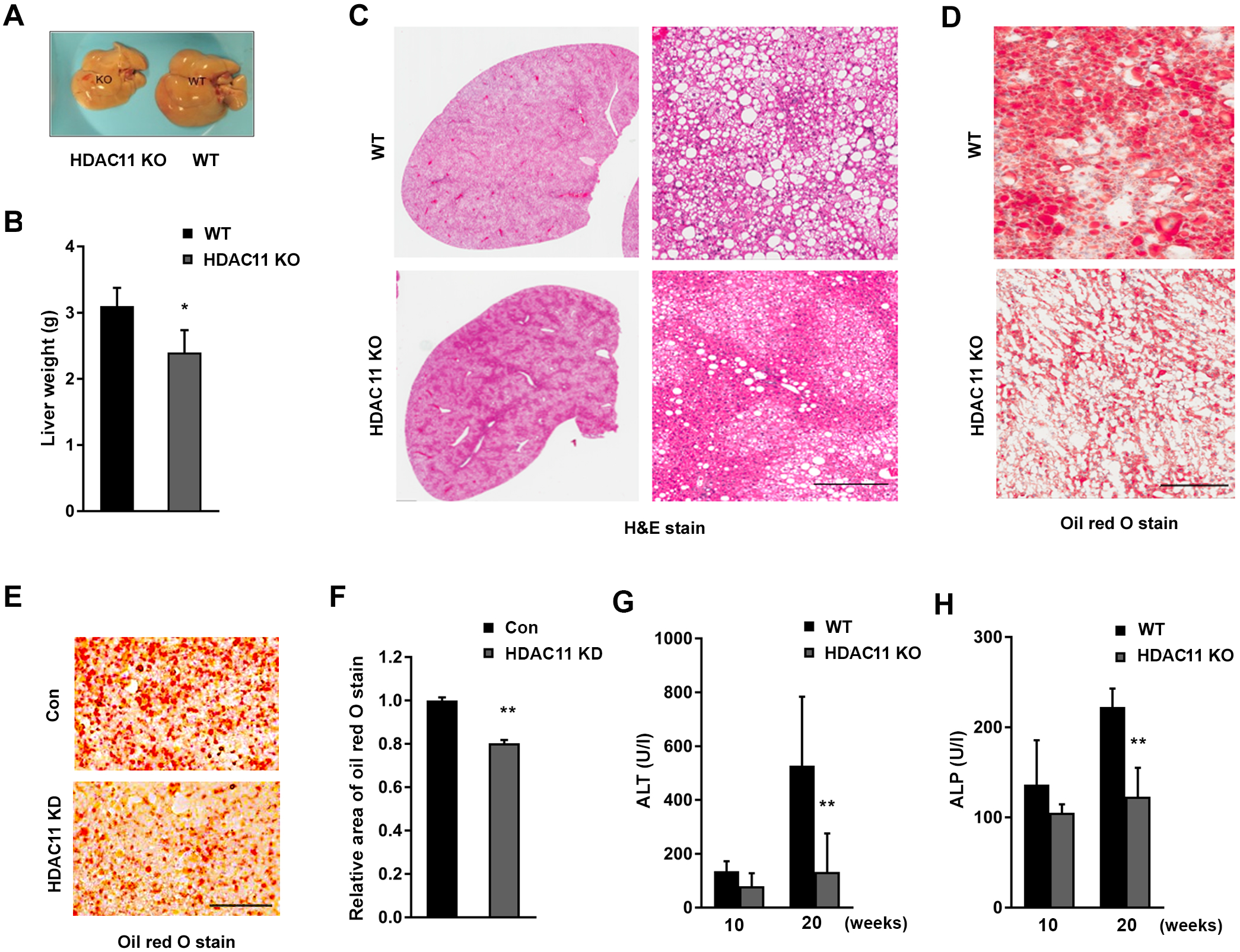
HDAC11 KO mice are protected from HFD-induced hepatic steatosis. (A) Representative pictures of liver morphology and (B) liver weight from WT (n=7) and HDAC11 KO (n=6) mice on 26-week HFD. (C) Representative H & E stain and (D) oil red O stain of liver sections from WT and HDAC11 KO mice on 26-week HFD, scale bar: 300μm. (E) Representative images of oil red O stain and (F) relative area of oil red O stain quantified from AML-12 control (Con) cells and HDAC11 knockdown (HDAC11 KD) cells after 5 days of free fatty acid treatment, scale bar: 100μm. (G) ALT and (H) ALP plasma levels between WT (n=8) and HDAC11 KO (n=10) mice on 10-week and 20-week HFD. Data shown are mean ±SD. *p<0.05, **p<0.01.

The levels of serum alanine transaminase (ALT, an indicator of liver damage) and alkaline phosphatase (ALP, a marker of bile duct obstruction, intrahepatic cholestasis, or liver damage) were measured, and HDAC11 KO mice completely blocked the HFD-elicited elevation of ALT (Fig. 3G) and ALP (Fig. 3H). However, there is no significant difference in albumin (Fig. S4A) or total bilirubin levels (Fig. S4B) between HDAC11 KO and WT mice.

### HDAC11 deficiency enhances thermogenic potential and promotes UCP1 expression

Total energy expenditure in mammals is a sum of energy utilization during external physical activity and internal heat production. After monitoring locomotor activity with the Phenomaster System for four consecutive days, the total locomotor activity in the XYZ plane, horizontal activity in the XY plane, and rearing activity in the Z plane of WT and HDAC11 KO mice, were compared and showed no statistical differences (Fig S5A-H). These results argue that the resistance of HDAC11 KO mice to HFD-induced obesity depends more on metabolic mechanisms than on physical activity.

Defect in non-shivering thermogenesis appears to be constitutive, rather than a secondary consequence of obesity (16). We found that body temperature was significantly increased in both normal (ND)-and HFD-fed HDAC11 KO mice compared to WT mice at room temperature (Fig. 4A). By performing the cold challenge experiment (4°C for 6-hour), in stark contrast to WT mice, HDAC11 KO mice under either ND or HFD exhibited a marked increase in thermogenesis and effective resistance to a drop in body temperature (Fig. 4B).

**Figure 4.**
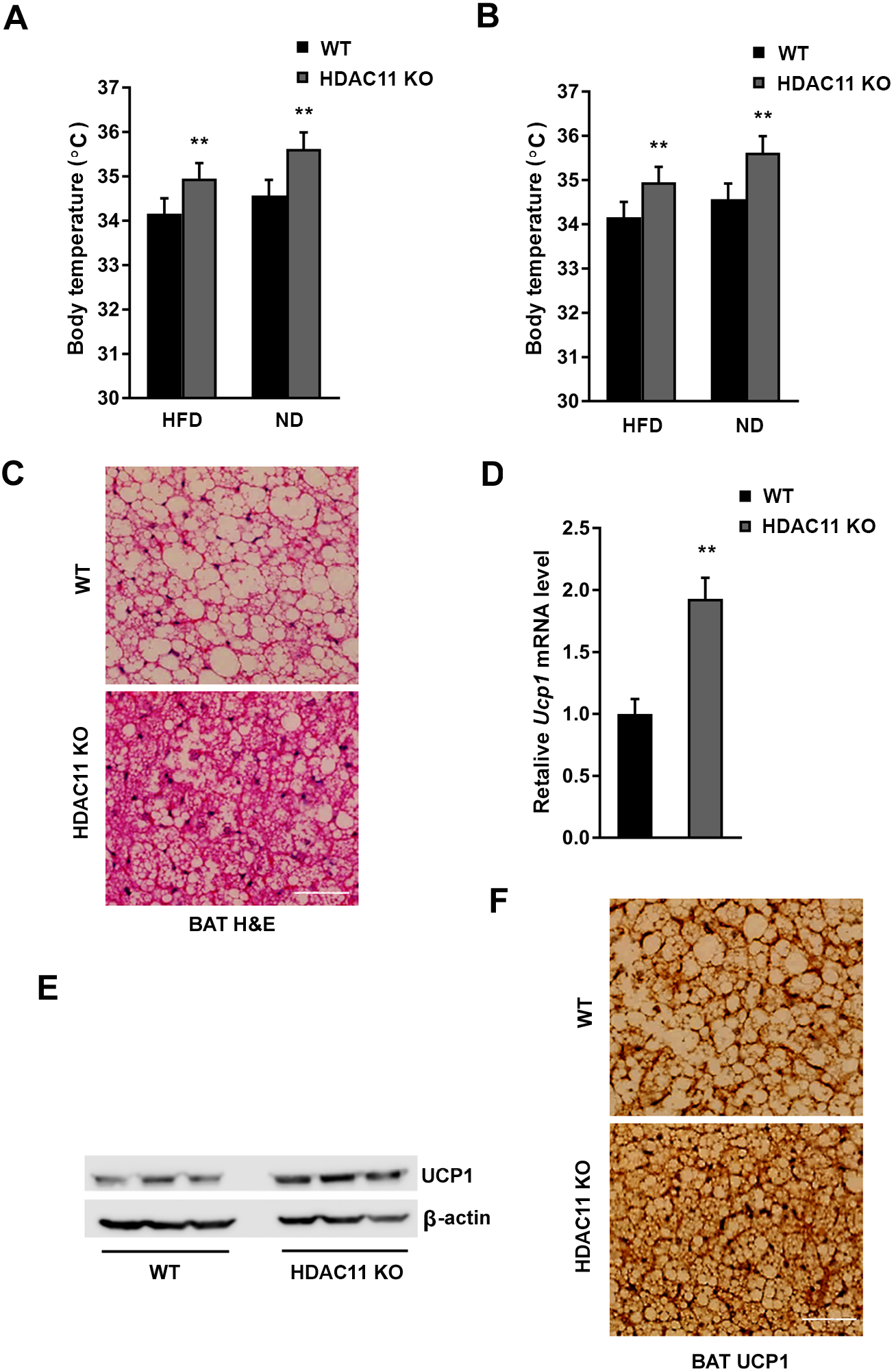
HDAC11 deficiency promotes thermogenesis in mice. (A) Body temperatures of mice on HFD (WT, n=8; HDAC11 KO, n=8) and ND (WT, n=7; HDAC11 KO, n=5). (B) Body temperatures after 4°C cold exposure for 6 h on HFD (WT, n=8; HDAC11 KO, n=8) and ND (WT, n=7; HDAC11 KO, n=5). (C) Representative H & E stain of brown adipose tissue (BAT) in WT and HDAC11 KO mice on 13-week HFD, scale bar: 200μm. (D) Relative Ucp1 mRNA level in BAT of WT and HDAC11 KO mice on 13-week HFD. (E) Western blot analysis of UCP1 and β-actin protein level in BAT between WT (n=3) and HDAC11 KO (n=3) mice. (F) Representative images of UCP1 immunohistochemistry in BAT between WT and HDAC11 KO mice on 13-week HFD, scale bar: 200μm. Data shown are mean ±SD. *p<0.05, **p<0.01.

Brown adipose tissue (BAT) is specialized in adaptive thermogenesis through heat production in response to cold or excess calories. It was unequivocally demonstrated that the majority of human adults possess active BAT, which is associated with a favorable metabolic phenotype (17). As shown in Fig. 4C, H & E staining of BAT demonstrates a condensed tissue structure with significantly smaller lipid droplets in HDAC11 KO mice than in WT mice. UCP1 is responsible for non-shivering thermogenesis in BAT. Mice deficient in UCP1 have been shown to be susceptible to weight gain, whereas overexpression of UCP1 provides protection against diet-induced obesity (18, 19). Our results indicate a marked increase in Ucp1 mRNA (Fig. 4D) and protein (Fig. 4E) expressions in the BAT of HDAC11 KO mice under 13 weeks of HFD. The immunohistological analysis revealed a diffused distribution of UCP1 staining at a slightly higher intensity in the BAT of HDAC11 KO mice (Fig. 4F).

### HDAC11 deficiency promotes oxygen consumption and elevates CPT1 activity

Oxygen consumption has been used as the golden standard for indirect calorimetry. To further understand the resistance to HFD-induced obesity in HDAC11 KO mice, we measured oxygen consumption (VO2), carbon dioxide production (VCO2) and respiratory exchange ratio (RER; VCO2/VO2). As shown in Figs. 5A, B, the 24-hour oxygen consumption and carbon dioxide production of HDAC11 KO mice are significantly elevated across both light and dark phases. As expected, there was no difference in RER level between WT and HDAC11 KO mice. However, an increased RER of HDAC11 KO mice in the dark phase was statistically significant, when compared to that in the light phase (Fig. 5C), suggesting that HDAC11 KO reversed the HFD-induced disruption of circadian rhythms. The energy expenditure could also be assessed through the metabolic rate, which is calculated with the parameters of VO2, VCO2 and body weight. As shown in Fig. 5D, deletion of HDAC11 resulted in a significant elevation of total calories expenditure across light and dark phases, highlighting the importance of enhanced metabolic activity in HDAC11 KO mice for resistant to obesity.

**Figure 5.**
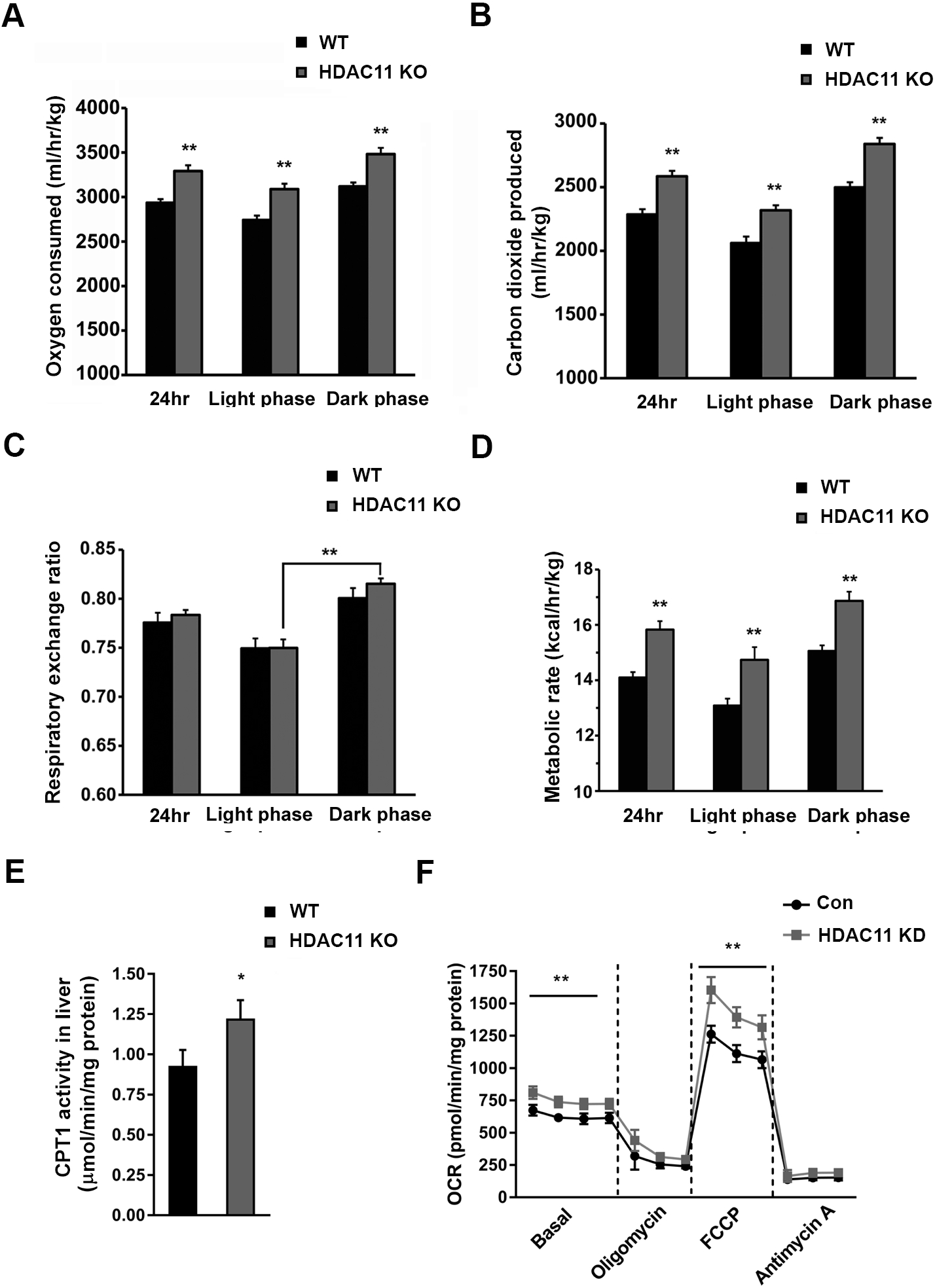
Lack of HDAC11 promotes oxygen consumption and elevates CPT1 activity. (A) Oxygen consumed, (B) carbon dioxide produced, (C) respiratory exchange ratio and (D) metabolic rate of WT (n=4) and HDAC11 KO (n=4) mice in 24 hours, light phase and dark phase respectively. (E) Quantitative CPT1 activity in liver tissues between WT (n=4) and HDAC11 KO (n=4) mice. (F) Oxygen consumption rate (OCR) in control (Con) and HDAC11 knockdown (HDAC11 KD) AML-12 cells in basal conditions, or in the presence of 1μM oligomycin, 1μM FCCP, 0.5μM Antimycin A. Data shown are mean ±SD. *p<0.05, **p<0.01.

Given our observation that the absence of HDAC11 promoted oxygen consumption and metabolic activity, we hypothesize that fat oxidation serves as a crucial mechanism for energy expenditure. Carnitine palmitoyltransferase 1 (CPT1), which initiates the first step of the oxidative pathway in the mitochondrial matrix, is the key regulatory enzyme of mitochondrial long-chain fatty acid β-oxidation (20). As shown in Fig. 5E, liver CPT1 activity is markedly elevated in HDAC11 KO mice.

To further verify the impact of HDAC11 deficiency on mitochondrial respiration, we measured the mitochondrial oxygen consumption rate (OCR) under basal and stress conditions in AML12 cells. In comparison to control cells, HDAC11 KD cells showed an increased OCR even under basal conditions (Fig. 5F). HDAC11 KD also significantly elevated maximal respiration driven by a mitochondrial uncoupling agent, carbonyl cyanide-4-(trifluoromethoxy) phenylhydrazone (FCCP). Moreover, the spare respiratory capacity (difference between maximal and basal respiration) was markedly elevated in HDAC11 KD cells, suggesting that a decrease in HDAC11 enhances the ability to respond to an increased energy demand under stress conditions (21).

### HDAC11 deficiency elevates plasma adiponectin and activates the adipoR-AMPK signaling pathway in the liver

Adiponectin, the most abundant adipose-specific adipokine, is a hormone involved in regulating glucose levels and fatty acid degradation. An association between reduced adiponectin production and the pathogenesis of non-alcoholic fatty liver disease (NAFLD) has been confirmed in numerous studies (22, 23). Results from our ELISA analysis revealed significantly elevated plasma adiponectin levels in 20-week HFD-fed HDAC11 KO mice (Fig. 6A). The *AdipoQ* gene mRNA expressions in the eWAT, BAT, and liver of HDAC11 KO mice were also markedly increased compared to the WT animals (Fig. 6B).

**Figure 6.**
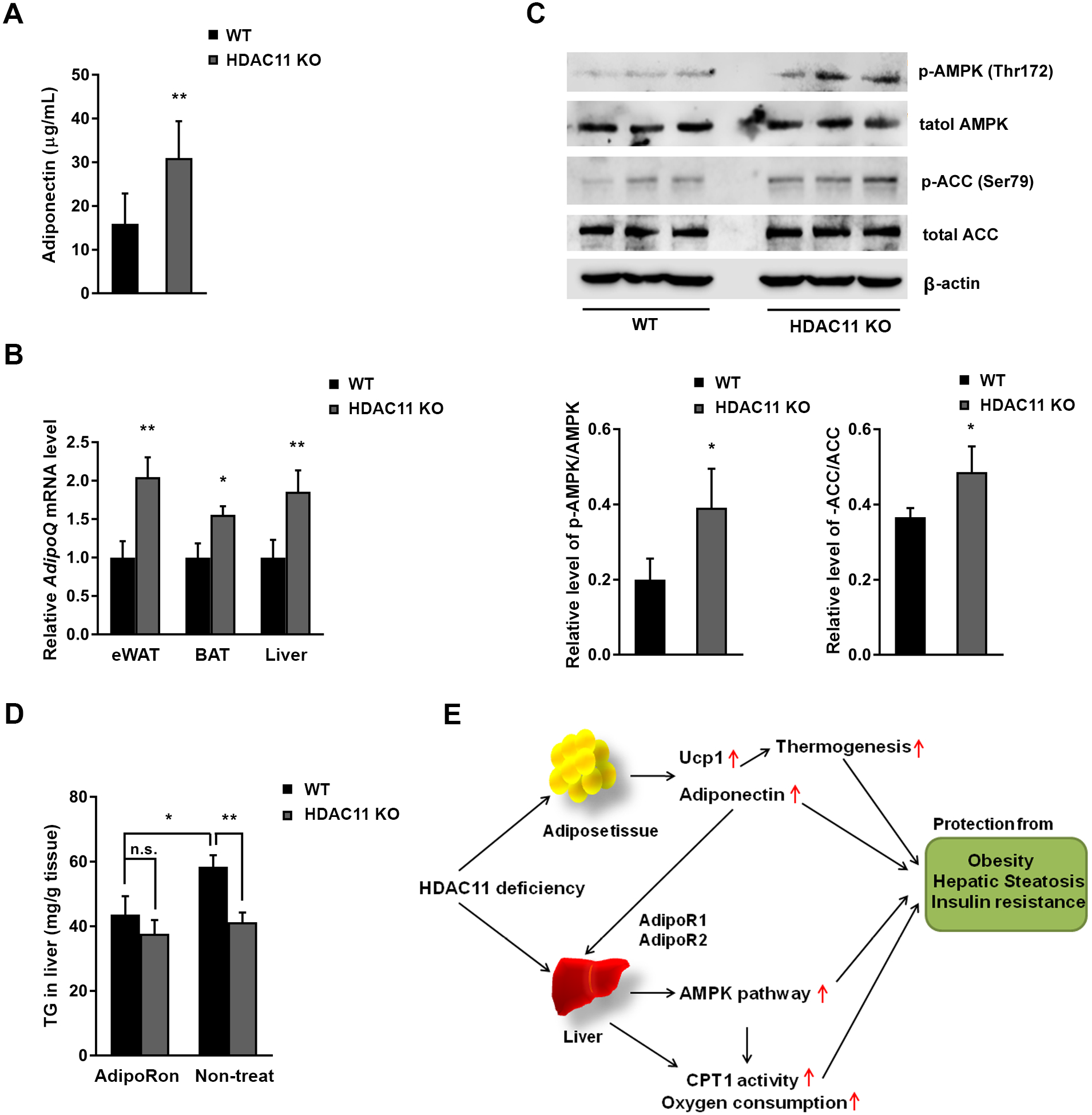
HDAC11 deficiency elevates plasma adiponectin and activates AdipoR-AMPK signaling in the liver. (A) Plasma adiponectin level in WT (n=5) and HDAC11 KO (n=6) mice on 20-week HFD. (B) Relative AdipoQ mRNA expression levels in eWAT, BAT and liver tissue between WT (n=3) and HDAC11 KO (n=3) mice. (C) Western blot analysis of liver p-Thr172-AMPK, AMPK, p-Ser79-ACC, ACC, and β-actin protein levels in WT (n=3) and HDAC11 KO (n=3) mice on 20-week HFD. Lower panel shows relative ratios of p-AMPK/AMPK and p-ACC/ACC. (D) Hepatic triglyceride (TG) content in WT and HDAC11 KO mice fed HFD for 30 days, then 10 days of AdipoRon (50mg/kg body weight) treatment (WT, n=5; HDAC11 KO, n=4) or non-treatment (WT, n=5; HDAC11 KO, n=4). (E) A schematic diagram of HDAC11 deficiency and its effect on metabolic syndrome in HFD mice. Data shown are mean ±SD. *p<0.05, **p<0.01.

Adiponectin signaling is mainly mediated through the AdipoR1-AMP-activated protein kinase (AMPK) and AdipoR2-peroxisome proliferator-activated receptor α (PPARα) pathways in the liver (24). Consistent with previous reports that AdipoR2 mRNA levels reduced in NAFLD patients and hepatocytes induced by fatty acid (25), RT-qPCR analysis confirmed an increase in adiponectin receptor 2 (AdipoR2), but not adiponectin receptor 1 (AdipoR1), in the livers of HDAC11 KO mice (Fig. S6).

AMPK signaling plays an essential role in adiponectin-mediated beneficial effects on adipose tissue and lipid metabolism. One of the most well-defined mechanisms of AMPK activation is phosphorylation at Thr172 of the α-subunit (26). As shown in Fig. 6C, compared to WT animals, Thr172 of AMPK is hyperphosphorylated in hepatic tissues from HDAC11 KO mice. Additionally, phosphorylation at Ser79 of acetyl-CoA carboxylase (ACC), a downstream substrate of AMPK (27), was increased in parallel with AMPK activity (Fig. 6C). These results demonstrate that HDAC11 deletion increases AMPK activity and stimulates the AMPK-ACC signaling pathway.

Previous studies using AdipoR1 KO mice showed that the loss of AdipoR1 increased adiposity associated with decreased glucose tolerance and energy expenditure. However, AdipoR2 KO mice were lean and resistant to HFD-induced obesity associated with improved glucose tolerance, higher energy expenditure, and reduced plasma cholesterol levels (28). To further study the effect of HDAC11 and adiponectin, we activated AdipoR with AdipoRon, an adiponectin receptor agonist that binds both AdipoR1 and AdipoR2 (29), in AML12 cells. AdipoRon activated the phosphorylation of Thr172-AMPK and Ser79-ACC in a dose-dependent manner in both control and HDAC11 KD AML12 cells (Fig. S7A). Oral administration of HFD-fed mice with AdipoRon (50mg kg^-1^ body weight/day) for 10 days revealed no further increase of phospho-AMPK and phospho-ACC in the liver of HDAC11 KO mice, suggesting a maximized signaling stimulation by AdipoRon. (Fig. S7B). Indeed, AdipoRon treatment effectively prevents an increase in liver TG in HFD-fed WT mice, and because the receptor agonist did not further inhibit liver TG levels in HDAC11 KO mice, it is reasonable to conclude that HDAC11 deficiency-activated adiponectin-AdipoR-AMPK pathway serves as a major mechanism in the attenuation of hepatic TG accumulation (Fig. 6D).

In summary, our results convincingly show that HDAC11 is an important metabolic regulator. Mice that lack HDAC11 are protected from diet-induced obesity and fatty liver. HDAC11 deletion inhibited HFD associated hypercholesterolemia, hyperinsulinemia and reversed liver injury. HDAC11 deficiency boosted thermogenesis, which is associated with increased UCP1 expression in BAT. The loss of HDAC11 elevated CPT1 activity and mitochondrial oxygen consumption, a key pathway for energy expenditure. A significant increase in adiponectin production by HDAC11 deletion might serve as another important regulatory mechanism that activates adiponectin-associated signaling and re-programming (Fig. 6E).

## Discussion

In this study, we show that the absence of HDAC11 prevented obesity and ameliorated pathological manifestations associated with chronic high-fat feeding in mice, including glucose intolerance, insulin resistance, hypercholesterolemia, and hepatosteatosis. Under HFD, compared to WT animals, HDAC11 KO mice were healthy and showed no apparent abnormality in their appearances or behaviors throughout our studies.

Accumulating evidence indicates a link between metabolic disorders and dysregulated HDACs. For example, in sharp contrast to HDAC11 KO mice, liver-specific HDAC3 KO resulted in induced obesity and hepatosteatosis accompanied with hepatocyte hypertrophy, liver damage with elevated ALT, and a significant increase in serum triglyceride, cholesterol, and LDL (3, 4, 30). These results suggest that HDAC11 reprograms metabolism through novel and unique signaling pathways, and each HDAC may have distinct functions in metabolic regulations. Thus, our study illuminates the potential therapeutic benefit of selective HDAC11 inhibitors for the prevention or treatment of obesity and obesity-related diseases.

By catalyzing histone deacetylation, HDACs are thought to modify chromatin and regulate gene transcription. However, recent studies suggest that many HDACs possess enzymatic activities in addition to deacetylation and most, if not all, HDACs have additional nonhistone substrates and function beyond transcriptional regulation. Although HDAC11 contains conserved catalytic core regions shared with class I and II HDACs, we have not been able to detect significant HDAC11 deacetylase activity in HDAC11 overexpressing cells (Figs. S8A, B). Our results are consistent with a recent report that HDAC11 is a fatty-acid deacylase rather than deacetylase, and this function of HDAC11 is phylogenetically conserved (14). Identification of HDAC11 de-fatty-acylation targets will help to further understand how the lack of HDAC11 prevents obesity in animals as well as the roles and functions of HDAC11-mediated metabolic reprogramming.

BAT is a regulator of energy expenditure and body fat in humans and rodents. We found an increase in thermogenesis in HDAC11 KO mice accompanied with high expression of UCP1, which is responsible for uncoupling and proton leak across the inner mitochondrial membrane and heat production. Increased BAT capacity of thermogenesis relies on the expression of UCP1, which most likely contributes to the HDAC11 KO phenotypes under HFD in mice. While numerous transcriptional regulatory mechanisms associated with UCP1 expression have been proposed, it remains unclear if they are phylogenetically conserved and hence essential for UCP1 expression (31). HDAC3 has been reported to activate the *Ucp1* enhancer to ensure thermogenic aptitude (32). Mice with BAT-specific ablation of HDAC3 become severely hypothermic with nearly no UCP1 expression. Despite the similarity in the conserved catalytic core regions between HDAC3 and HDAC11, HDAC11 deletion promoted thermogenic function in BAT with an increased UCP1 expression. Currently, we do not yet know if the HDAC11 defatty-acylation activity is responsible for the underlying mechanism by which HDAC11 regulates UCP1 expression.

Previously, compelling evidence demonstrated that adiponectin enhances insulin sensitivity, improves fatty acid oxidation, and markedly ameliorates obesity-associated pathologic symptoms (33). The biological function of adiponectin is mediated, at least in part, by the existence of different oligomeric complexes of plasma adiponectin (trimers, hexamers, and high molecular weight multimers) (34). Although adiponectin is predominantly produced by adipose tissues, plasma adiponectin concentration and adiponectin gene expression are inversely correlated with adiposity. The regulatory roles of Class I and II HDACs in adiponectin expression may be different for each HDAC. Upregulation of HDAC9 expression blocked the adipogenic differentiation program under HFD animals, leading to an accumulation of improperly differentiated adipocytes with a diminished expression of adiponectin (9). Valproic acid, a class I HDAC inhibitor, significantly decreased adiponectin protein and mRNA levels in both mice and 3T3-L1 adipocytes (35). Sodium butyrate (NaB), a class I and IIa HDAC inhibitor, stimulated adipocyte differentiation and adipogenic gene expression, including adiponectin (36). Here, our data convincingly shows that the loss of HDAC11 not only elevates plasma adiponectin, but also promotes adiponectin gene expression in adipose tissue.

The importance of adiponectin in the balance of hepatic TG was demonstrated by quantitative analysis of hepatic TG in HFD-fed adiponectin KO mice, which showed a markedly elevated TG level which was reversed by adiponectin treatment (37). The targets of adiponectin include lysophospholipids, which are up-regulated by HFD and contribute to the phenotypes of HFD-induced insulin resistance, impaired glucose tolerance, and hypertriglyceridemia (37). The selective reduction of ceramides (C20:0 and C18:0) by adiponectin further suggests its influence on *de novo* and salvage pathways of ceramide synthesis (38). Ceramide is the neutral lipid building block of sphingolipids or, if they contain sugars, of glycosphingolipids. These lipids not only serve structural roles in biomembranes, but also have wide effects on signal transduction and cell function. Although the mechanisms whereby lipids exert their functions have been less clear than those of proteins, our findings on the important regulatory role of HDAC11 on adiponectin reveals a new physiological signaling pathway, which confers distinct biochemical properties. Furthermore, the recent discovery of HDAC11 as a fatty-acid deacylase provides a new opportunity to investigate this previously under-explored protein posttranslational modification and its impact on adiponectin cell signaling, membrane trafficking, protein-membrane interactions, and cellular localization.

AMPK signaling is crucial in maintaining energy homeostasis, and AMPK has been implicated in the regulation of a number of metabolic pathways in the liver, including gluconeogenesis, fatty acid synthesis, and fatty acid oxidation (39). Phosphorylation of Thr172 in AMPK, catalyzed by liver kinase B1 (LKB1), calcium/calmodulin-dependent kinase kinase 2 (CaMKK2) and TGFβ-activated kinase1 (TAK1), could activate AMPK. Active AMPK subsequently associates with phospho-Ser79 in acetyl-CoA carboxylase (ACC). Phospho-ACC could enhance the activity of CPT1, a rate limiting enzyme in fatty acid oxidation, thereby preventing excess TG storage in the liver (26, 40). We have provided evidence that HDAC11 impacts plasma adiponectin levels, adiponectin-AMPK activation, and adiponectin-AMPK signaling-mediated metabolism outcomes, such as hepatic TG accumulation.

In conclusion, our study reveals a novel physiological function for HDAC11, which regulates metabolism homeostasis through boosting UCP1 expression, promoting CPT1 enzyme activity, elevating plasma adiponectin, and activating adiponectin-AdipoR-AMPK pathway. These findings warrant further investigations into targeting HDAC11 for obesity and related metabolic diseases.

## Methods

### Animals

WT and *HDAC11* KO mice are on C57BL/6 background. To assess metabolic parameters, male HDAC11 KO and WT mice at 3 weeks of age were fed a HFD (Adjusted Calories Diet, TD. 88137, Harlan, 42% kcal from fat, 42.7% kcal from carbohydrate, and 15.2% kcal from protein) for the indicated times. Mice body temperature recordings were determined with YSI 4600 Precision thermometer (YSI, Inc., Yellow Springs, OH).

### Comprehensive Cage Monitoring System

Comprehensive metabolic cages were used to identify altered physiological systems in response to HFD. Phenomaster System (TSE Systems, USA) accommodating 12 mice simultaneously was used to measure calorimetry, activity, circadian patterns, water and food intake for 7 consecutive days. The equipment measured the volumes of O2 consumed and CO2 produced in air samples independently from each cage, calculating respiratory exchange ratio, and metabolic rate corrected for body weight. Food and water intake were measured directly and corrected for spillage. Locomotor activity was measured every 10 seconds by counting the number of infrared beam breaks in the X, Y, and Z planes. After a 3-day habituation period to the metabolic cages, 96 h of data was analyzed, with the reviewer blind to the genotype, for all measurements.

### Cell culture

Mouse normal hepatocyte AML12 was purchased from American Type Culture Collection. Cells were grown in a 1:1 mixture of Dulbecco’s modified Eagle’s medium and Ham’s F12 medium with 0.005 mg/ml insulin, 0.005 mg/ml transferrin, 5 ng/ml selenium and 40ng/ml dexamethasone with 10% fetal bovine serum (FBS), at 37°C in a humidified atmosphere of 5% CO2.

### Antibodies and reagents

AMPKα (2532), phosphor-AMPKα (T172) (2531), ACC (3662) and phosphor-ACC (S79) (11818) antibodies were purchased from Cell Signaling Technology. UCP1 (ab23841) antibody was purchased from Abcam. β-actin (A5316) antibody was purchased from Sigma-Aldrich. Mouse IgG (NA931V) and rabbit IgG (NA934V) HRP-linked whole antibodies were purchased from GE Healthcare. To generate an anti-HDAC11 antibody, a peptide corresponding to amino acids 179-193 of HDAC11 (DLDAHQGNGHERDFM) was injected subcutaneously into New Zealand white rabbits (Alpha Diagnostic, Inc.). The resulting antibody was immune-affinity purified on a peptide column with Affi-gel agarose (Bio-Rad). Specificity of this anti-HDAC11 antibody was confirmed using brain tissue extracts from HDAC11 KO and WT mice. Oil red O (O0625) and oleic acid (O3008) were purchased from Sigma-Aldrich. AdipoRon (15941) was purchased from Cayman Chemical, and Insulin-Transferrin-Selenium (41400-045) was from Thermo Fisher Scientific.

### Histology and Immunohistochemistry

Mice were anesthetized with ketamine-xylazine solution and perfused intracardially with PBS and 4% formalin solution through the left ventricle. Liver, brown and white adipose tissue were dissected and fixed in 4% phosphate-buffered formalin solution at room temperature overnight, and embedded in paraffin wax. Paraffin sections (6 μm) were cut and stained by hematoxylin and eosin (H & E). For immunohistochemistry, tissue paraffin sections were deparaffinized and rehydrated by immersing in xylene and ethanol, followed with antigen retrieval using the citrate buffer method. Sections were incubated in 3% H2O2 solution at room temperature for 15 min, and blocked by 5% donkey serum in PBS-0.1% tween-20 for 1 hour. Next, sections were incubated with primary antibody overnight at 4°C followed by incubation with biotinylated secondary antibody (Vector Laboratories, MP-7500) for 30 min at room temperature. DAB substrate solution (Vector Laboratories, SK-4105) was added to the sections according to the manufacturer’s instructions.

### Blood analyses

Plasma total cholesterol, TG, glucose, ALT, ALP, Albumin and total bilirubin levels were measured using FUJI DRI-CHEM slide live panel (FUJIFILM), and were evaluated by DRI-CHEM 4000 chemistry analyzer. Hematology results were measured by HemaTrue Veterinary Chemistry Analyzer.

### Glucose and Insulin tolerance test

Mice were fasted 12 hours before the administration of glucose or insulin. Before the test, mice were tail nicked at the very distal tip portion to collect 1-2 μl of blood to measure baseline glucose levels on a portable glucometer. Glucose (1 mg/gram body weight) or insulin (1 mU/gram body weight) was injected intraperitoneal and glucose levels were tested at 15, 30, 60, 90 minutes after injection until glucose returned to baseline level.

### Triglyceride (TG), Insulin and Adiponectin quantification

Liver TG levels were measured by Triglyceride Colorimetric Assay Kit (Cayman Chemical, Catalog #10010303) following the procedure provided by the manufacturer. Insulin and adiponectin in the plasma were tested by an enzyme-linked immuno sorbent assay (Insulin mouse ELISA kit was from Thermo Fisher Scientific, Catalog #EMINS, and adiponectin mouse ELISA kit was from Crystal Chem, Catalog #80569).

### Oil Red O stain

Cryosections of livers were dried for 30-60 minutes at room temperature and then fixed in ice cold 10% formalin for 5-10 minutes. Sections were rinsed immediately in 3 changes of distilled water, then in absolute propylene glycol for 2-5 minutes. Slides were stained in 0.5% oil red O solution for 10 minutes in 60°C oven, then differentiated in 85% propylene glycol solution for 5 minutes. Slides were washed with distilled water and mounted with glycerin jelly.

For AML12 cells, control and HDAC11 KD cells were cultured for 5 days in complete medium with free fatty acid (200 μM oleic acid, Sigma, O1383). To detect lipid accumulation, cells were stained with oil red O. Cells were washed twice with PBS, fixed in 4% paraformaldehyde for 1 hour, stained with 0.3% Oil Red O in 60% isopropanol for 10 min, and washed with 60% isopropanol. Then cells were washed with PBS and photographed.

### Immunoblotting

Cells were lysed in NETN buffer (10 mM Tris, pH 8.0, 150 mM NaCl, 1 mM EDTA, 1% NP-40, and protease inhibitor cocktail) with 2 mM sodium orthovanadate, 2 mM sodium fluoride, 50mM nicotinamide and 25 mM β-glycerophosphate. Fresh mice tissues were quickly frozen with liquid nitrogen, and homogenized in NETN buffer with 2 mM sodium orthovanadate, 2 mM sodium fluoride, 50mM nicotinamide and 25 mM β-glycerophosphate in a Potter-Elvehjem homogenizer followed by ultrasonication on ice. Samples were centrifuged for 10 min at 12,000g at 4°C, and protein contents of the samples were determined by the Bradford assay. Proteins were separated by SDS-PAGE and transferred to polyvinylidene fluoride (PVDF) membrane (Millipore, MA), which was probed with primary and secondary antibodies. Proteins were visualized using a chemiluminescence detection kit (Pierce).

### CPT1 activity

The non-radioactive CPT1 activity assay is used with Ellman’s reagent (41). A yellow-colored product with the release of CoA-SH from palmitoyl-CoA, is measured at an absorbance of 412 nm using the 5, 5’-dithio-bis-(2-nitrobenzoic acid) (DTNB, Sigma, D8130). Mice liver whole protein lysates were prepared as described for immunoblotting. Whole protein samples (30 μg) and DTNB (final concentration 1 mM) were mixed in the reaction buffer (20mM Tris, pH 8.0, 1mM EDTA) and incubated for 30 min at room temperature. To start the reaction, palmitoyl-CoA (Pubchem, 644109) (100 μM final concentration) and L-carnitine (Sigma, C0283) (5 mM final concentration) solutions were added to the reaction system. As a negative control, reaction buffer was used instead of palmitoyl-CoA. Absorbance was recorded at 412 nm at 1 min intervals for 90 min. Activity was defined as nmol CoA-SH released/min/mg protein.

### Oxygen Consumption Rate Measurements

Oxygen consumption rate (OCR) of AML12 cells was analyzed by XF-96 Extracellular Flux Analyzer (Seahorse Bioscience). Cells were washed with XF assay medium and placed into a 37 °C incubator without CO2 for 45 minutes to 1 h. For the mitochondrial stress test, the cells were treated with the ATP synthase inhibitor oligomycin (1.0 μM), the chemical uncoupler FCCP (1.0 μM), and the electron transport inhibitor antimycin A (0.5 μM). Basal oxygen consumption was assessed before the addition of any mitochondrial inhibitor. OCR values were normalized to cell number.

### Preparation and transduction of shRNA

The lentivirus transduction particles containing shRNA, specific for mouse HDAC11 (SHVRS-NM_144919) or non-targeting shRNA (SHC002V), were purchased from Sigma-Aldrich. Lentivirus was prepared by transfecting into HEK 293T cells together with packaging vectors. Packaged lentivirus was transfected into AML12 cells with polybrene, and transduced cells were selected for resistance to puromycin.

### RT-qPCR

Total RNA was isolated from mouse tissues using the mirVana isolation kit (Applied Biosystems). Total RNAs from at least three mice were mixed equally. Reverse transcription was carried out by the qScript cDNA synthesis kit (Quanta Biosciences, Gaithersburg, MD). Relative quantitation of mRNAs was tested via SYBR green-based quantitative PCR and specific primers (Table S2). qPCR was carried out by 7900 HT fast real-time PCR system (Applied Biosystems) following the manufacturer’s instruction of iQ SYBR green Supermix (BioRad). Mice *GAPDH* served as an internal control, and PCR results were analyzed using the 2^-(ΔΔCΓ)^ method.

### Statistical analysis

Statistical differences were detected using t-tests when only comparing two groups or ANOVA plus post-hoc comparison for multiple comparisons with significance set at p<0.05, and very significance set at p<0.01. (GraphPad Prism 7.03).

## Acknowledgements

We thank Thomas Rosahl at Merck for the *Hdac11* constitutive knockout mice, which was generated at the Institut Clinique de la Souris and obtained via Taconic. We also thank Tim McKinsey and Rushita Bagchi for insightful discussions on the project. Additionally, we thank Raneen Rahhal and Sonali Bahl for critical reading of the manuscript. This project was supported by grants from the NIH.

## Author contributions

E.S. planned and supervised the research. L.S., C.M.D., A.A., and E.T. carried out the experiments. L.S., C.M.D., and E.S. analyzed the data. L.S. C.M.D., K.B., and E.S. wrote the manuscript.

## References

1. Seto E & Yoshida M (2014) Erasers of histone acetylation: the histone deacetylase enzymes. Cold Spring Harb Perspect Biol 6(4):a018713.

2. Li F, et al. (2016) Histone Deacetylase 1 (HDAC1) Negatively Regulates Thermogenic Program in Brown Adipocytes via Coordinated Regulation of Histone H3 Lysine 27 (H3K27) Deacetylation and Methylation. The Journal of biological chemistry 291(9):4523–4536.

3. Sun Z, et al. (2012) Hepatic Hdac3 promotes gluconeogenesis by repressing lipid synthesis and sequestration. Nat Med 18(6):934–942.

4. Knutson SK, et al. (2008) Liver-specific deletion of histone deacetylase 3 disrupts metabolic transcriptional networks. EMBO J 27(7):1017–1028.

5. Lundh M, Galbo T, Poulsen SS, & Mandrup-Poulsen T (2015) Histone deacetylase 3 inhibition improves glycaemia and insulin secretion in obese diabetic rats. Diabetes Obes Metab 17(7):703–707.

6. Ferrari A, et al. (2017) Attenuation of diet-induced obesity and induction of white fat browning with a chemical inhibitor of histone deacetylases. Int J Obes (Lond) 41 (2):289–298.

7. Tian Y, et al. (2015) Histone Deacetylase HDAC8 Promotes Insulin Resistance and beta-Catenin Activation in NAFLD-Associated Hepatocellular Carcinoma. Cancer Res 75(22):4803–4816.

8. Bricambert J, et al. (2016) Impaired histone deacetylases 5 and 6 expression mimics the effects of obesity and hypoxia on adipocyte function. Mol Metab 5(12):1200–1207.

9. Chatterjee TK, et al. (2014) Role of histone deacetylase 9 in regulating adipogenic differentiation and high fat diet-induced metabolic disease. Adipocyte 3(4):333–338.

10. Purushotham A, et al. (2009) Hepatocyte-specific deletion of SIRT1 alters fatty acid metabolism and results in hepatic steatosis and inflammation. Cell Metab 9(4):327–338.

11. Xiao C, et al. (2010) SIRT6 deficiency results in severe hypoglycemia by enhancing both basal and insulin-stimulated glucose uptake in mice. The Journal of biological chemistry 285(47):36776–36784.

12. Schwer B, et al. (2010) Neural sirtuin 6 (Sirt6) ablation attenuates somatic growth and causes obesity. Proc Natl Acad Sci U S A 107(50):21790–21794.

13. Kanfi Y, et al. (2008) Regulation of SIRT6 protein levels by nutrient availability. FEBS Lett 582(5):543–548.

14. Kutil Z, et al. (2018) Histone Deacetylase 11 Is a Fatty-Acid Deacylase. ACS Chem Biol 13(3):685–693.

15. Biddinger SB, et al. (2005) Effects of diet and genetic background on sterol regulatory element-binding protein-1c, stearoyl-CoA desaturase 1, and the development of the metabolic syndrome. Diabetes 54(5):1314–1323.

16. Jung RT, Shetty PS, James WP, Barrand MA, & Callingham BA (1979) Reduced thermogenesis in obesity. Nature 279(5711):322–323.

17. Cypess AM, et al. (2009) Identification and importance of brown adipose tissue in adult humans. The New England journal of medicine 360(15):1509–1517.

18. Kopecky J, Clarke G, Enerback S, Spiegelman B, & Kozak LP (1995) Expression of the mitochondrial uncoupling protein gene from the aP2 gene promoter prevents genetic obesity. J Clin Invest 96(6):2914–2923.

19. Li B, et al. (2000) Skeletal muscle respiratory uncoupling prevents diet-induced obesity and insulin resistance in mice. Nat Med 6(10):1115–1120.

20. Jogl G, Hsiao YS, & Tong L (2004) Structure and function of carnitine acyltransferases. Ann N Y Acad Sci 1033:17–29.

21. Wagner BA, Venkataraman S, & Buettner GR (2011) The rate of oxygen utilization by cells. Free Radic Biol Med 51(3):700–712.

22. Polyzos SA, Kountouras J, & Mantzoros CS (2016) Adipokines in nonalcoholic fatty liver disease. Metabolism 65(8):1062–1079.

23. Adolph TE, Grander C, Grabherr F, & Tilg H (2017) Adipokines and Non-Alcoholic Fatty Liver Disease: Multiple Interactions. Int J Mol Sci 18(8).

24. Yamauchi T, et al. (2007) Targeted disruption of AdipoR1 and AdipoR2 causes abrogation of adiponectin binding and metabolic actions. Nat Med 13(3):332–339.

25. Shimizu A, et al. (2007) Regulation of adiponectin receptor expression in human liver and a hepatocyte cell line. Metabolism 56(11):1478–1485.

26. Kim J, Yang G, Kim Y, Kim J, & Ha J (2016) AMPK activators: mechanisms of action and physiological activities. Exp Mol Med 48:e224.

27. Fullerton MD, et al. (2013) Single phosphorylation sites in Acc1 and Acc2 regulate lipid homeostasis and the insulin-sensitizing effects of metformin. Nat Med 19(12):1649–1654.

28. Bjursell M, et al. (2007) Opposing effects of adiponectin receptors 1 and 2 on energy metabolism. Diabetes 56(3):583–593.

29. Okada-Iwabu M, et al. (2013) A small-molecule AdipoR agonist for type 2 diabetes and short life in obesity. Nature 503(7477):493–499.

30. Feng D, et al. (2011) A circadian rhythm orchestrated by histone deacetylase 3 controls hepatic lipid metabolism. Science 331 (6022):1315–1319.

31. Fromme T (2017) Commentary: Evolution of UCP1 Transcriptional Regulatory Elements Across the Mammalian Phylogeny. Front Physiol 8:978.

32. Emmett MJ, et al. (2017) Histone deacetylase 3 prepares brown adipose tissue for acute thermogenic challenge. Nature 546(7659):544–548.

33. Yamauchi T, et al. (2002) Adiponectin stimulates glucose utilization and fatty-acid oxidation by activating AMP-activated protein kinase. Nat Med 8(11):1288–1295.

34. Tsao TS, et al. (2003) Role of disulfide bonds in Acrp30/adiponectin structure and signaling specificity. Different oligomers activate different signal transduction pathways. The Journal of biological chemistry 278(50):50810–50817.

35. Qiao L, Schaack J, & Shao J (2006) Suppression of adiponectin gene expression by histone deacetylase inhibitor valproic acid. Endocrinology 147(2):865–874.

36. Yoo EJ, Chung JJ, Choe SS, Kim KH, & Kim JB (2006) Down-regulation of histone deacetylases stimulates adipocyte differentiation. The Journal of biological chemistry 281(10):6608–6615.

37. Liu Y, et al. (2015) Metabolomic profiling in liver of adiponectin-knockout mice uncovers lysophospholipid metabolism as an important target of adiponectin action. Biochem J 469(1):71–82.

38. Grosch S, Schiffmann S, & Geisslinger G (2012) Chain length-specific properties of ceramides. Prog Lipid Res 51 (1):50–62.

39. Steinberg GR & Kemp BE (2009) AMPK in Health and Disease. Physiol Rev 89(3):1025–1078.

40. Schreurs M, Kuipers F, & van der Leij FR (2010) Regulatory enzymes of mitochondrial beta-oxidation as targets for treatment of the metabolic syndrome. Obes Rev 11 (5):380–388.

41. Bieber LL, Abraham T, & Helmrath T (1972) A rapid spectrophotometric assay for carnitine palmitoyltransferase. Anal Biochem 50(2):509–518.

